# SPEx: Compartment-Resolved Proteomics via Expansion Microscopy–Guided Microdissection

**DOI:** 10.64898/2026.03.28.714993

**Authors:** Curdin A. Franziscus, Alexia Ferrand, Oliver Biehlmaier, Alexander Schmidt, Anne Spang

**Author notes:** Corresponding Authors: Anne Spang, Biozentrum, University of Basel, Spitalstrasse 41, CH-4056 Basel, Switzerland, Alexander Schmidt, Biozentrum, Biozentrum, University of Basel, Spitalstrasse 41, CH-4056 Basel, Switzerland.

## Abstract

Cells contain different organelles and compartments that are essential for cellular function and life. These organelles and compartments need to communicate to assess cellular state in a changing environment, adapt to the new situation, and to ensure functionality and homeostasis. Moreover, organization and communication differ between cell types. However, our knowledge about these changes is still rather scarce. Subcellular spatial proteomics aims to fill this knowledge gap. While proximity labeling techniques represent a great advance, they do not provide precise spatial resolution. To overcome this limitation, we developed SPEx (**S**ubcellular spatial **P**roteomics coupled to **Ex**pansion), in which we first expand cells about 10-fold, laser micro-dissect regions of interests and then perform mass spectrometry-based proteomics on these samples. We demonstrate the effectiveness of SPEx by determining the proteome of the Golgi, the nucleus and nucleoli. Importantly, we also identify novel components of these organelles. Combining inexpensive, already existing technologies makes SPEx readily usable by the wider scientific community.

## Introduction

Intracellular communication is essential for any life form. The communication, however, changes depending on a multitude of inputs such as extracellular cues, cell-cycle stage, nutrient availability, stress and age. This adaptability is key to the survival of any organism. While our understanding is increasing about intracellular communication in a few model organisms, such as tissue culture cells, Drosophila, *C. elegans* and yeast in unperturbed conditions, we still lack understanding how communication changes under different conditions and in different cell types. Efforts to fill this knowledge gap are hampered by fact that the basic machinery is often the same, but with variations in the interaction and composition of complexes, or specific temporal and spatial regulation.

Spatial subcellular proteomics intends to overcome this knowledge gap and aims to determine the localizations, concentrations and interactions of proteins within cells and tissues, while preserving their spatial information (Lundberg and Borner, 2019). A number of techniques have been developed to perform spatial proteomics at a subcellular resolution including differential centrifugation, organellar immunoprecipitation and proximity labeling methods (Fasimoye et al., 2023; Geladaki et al., 2019; Hundley et al., 2024; Lundberg and Borner, 2019; Roux et al., 2012). These techniques greatly improved our understanding of cellular organization and allowed the creation of comprehensive databases where large parts of, for example, the human proteome is mapped to specific subcellular compartments. However, all these approaches identify proteins based on proximity or co-purification with known markers and not on actual spatial positional information. This shortcoming cannot be circumvented since these techniques rely on a lysis step prior to proteomic analysis and hence all spatial information is inevitably lost.

More recently, Microscoop® aimed to overcome these drawbacks by introducing a technique that allows laser-guided biotinylation of specific subcellular regions *in situ*, whose protein content can be determined by subsequent cell lysis, biotin pulldown and liquid chromatography – mass spectrometry (LC-MS) based proteomic analysis (Chang et al., n.d.; Chen et al., 2023). While allowing for high spatial resolution, the laser-guided biotinylation of the region of interest (ROIs) is very time consuming and expensive. The subsequent affinity purification step requires the labeling of a rather large number of ROIs and cells, contributing to the substantial costs and low throughput. Because of the large number of required ROIs, this technique critically depends on automatization through machine learning algorithms and highly specialized equipment that are not readily available in most universities and research institutes. While an interesting technology, these drawbacks might limit the accessibility of Microscoop® for the wider scientific community.

Conversely, laser microdissection (LMD) allows direct LC-MS analysis of ROIs of tissues, cells or subcellular regions, and does not require a time- and sample-consuming affinity purification step. This method has been successfully applied to determine the proteome of single cells within tissues (Herrera et al., 2020; Large et al., 2020; Rosenberger et al., 2025, 2023) and is already an established and available workflow in many proteomics laboratories. Despite its recent success, this method is limited by its laser beam diameter (0.5 - 1 µm) which only allows collection of samples ranging from to 5-10 µm in diameter (Domazet et al., 2008; Walavalkar et al., 2025). The combination of expansion microscopy (ExM) and LC-MS has been applied in the past on tissues to isolate and analyze multi-cell regions within tissues with a diameter of ∼160 µm or more (Li et al., 2022). Technical limitations largely prevented its application for sub-cellular spatial proteomics apart from isolation of nuclei from 4x expanded cells (Dong et al., 2024a).

Here, we introduce SPEx (**S**ubcellular spatial **P**roteomics coupled to **Ex**pansion), a novel technique that extends the application of LMD and LC-MS analysis to single micron sub-cellular ROIs by combining 10x expansion of samples using a hydrogel with LMD. To allow for 10x expansion of cultured cells, SPEx utilizes the well-established TREx protocol developed for ExM (Damstra et al., 2022). This boost in expansion greatly increases the resolution of LMD and enables the dissection of individual organelles at a single micrometer resolution. Furthermore, we demonstrate that SPEx allows the determination of the proteome of a diverse group of organelles differing in size and shape. Moreover, we identified novel protein components of organelles. The quantitative aspect of our method provides also information on relative enrichment or depletion of proteins localizing to different parts/organelles in the cell. Strikingly, with the introduction of ultra-sensitive LC-MS platforms designed for single cell analysis (Brunner et al., 2022; Guzman et al., 2024), reasonable proteome coverage can be obtained with as little as about two dozen organelles. Moreover, SPEx is highly versatile and circumvents the need to develop specific protocols for the isolation of distinct organelles or cellular regions. SPEx relies exclusively on the visual selection of ROIs by the experimenter, based on fluorescent antibodies and/or dyes. In summary, we present SPEx, a novel kit-free, inexpensive, low-input method for *in situ* spatial proteomics at a micrometer resolution.

## Results

### Development of the SPEx method

#### Expansion Microscopy

To overcome the need for an accessible, inexpensive subcellular spatial proteomics method, we developed SPEx (Figure 1A). This method combines expansion microscopy, laser microdissection and LC/MS analysis. First, HeLa cells are fixed, and the cellular structure(s) of interest is/are labelled with antibodies as in standard immunofluorescence. In the present study, we labelled the Golgi apparatus with an anti-GM130 antibody. To visualize the cell, we labelled all proteins non-specifically with fluorescent N-hydroxysuccinimide (NHS) ester (Sheard et al., 2023). For the efficient dissection of subcellular structures, we aimed for a robust expansion of about 10x, while preserving downstream compatibility to LC/MS proteomic analysis. A protein disruption step is necessary to suppress the non-covalent interactions in the cells. Typically a protease treatment step (e.g. proteinase K), which is degrading proteins prior to expansion, is included to obtain high expansion factors (Chen et al., 2015; Tillberg et al., 2016; Wen et al., 2023). We cannot use protease treatment because it is incompatible with downstream LC-MS. Instead, we used SDS denaturation as described in the U-ExM or TREx protocols (Damstra et al., 2022; Gambarotto et al., 2019). To confirm that this alternative was suitable for LC/MS analysis, we tested whether such expanded samples still contained intact peptides that could be efficiently extracted from the gel. Indeed, the SDS treatment and expansion were compatible with proteomics. Moreover, we found that the sample size could be relatively small, while still maintaining good proteome coverage (Figure S1A and Table S1 and S2). While most of the ExM publications use protease treatment for disruption to reach the 10x expansion (Chen et al., 2015; Tillberg et al., 2016; Wen et al., 2023), SDS denaturation only gave us an expansion factor ranging from 4-7 fold, while the gels themselves expanded 10x. Due to this variability, we decided to adjust the TREx protocol (Damstra et al., 2022) to ensure a robust and high expansion factor. After reducing the anchoring between the proteins and matrix, and performing an overnight denaturation at 95 °C, we consistently obtained ∼9x expansion without protease treatment. This expansion represents an about 100x increase in area. At this expansion level, the fluorescence intensity was strong enough to detect the Golgi as well as proteins labelled with the NHS ester (Figure 1B and C). Of note, the strongest staining with the NHS ester was observed in the nucleus, and, in particular, in nucleoli. Because the nuclei were strongly labeled by the NHS ester, we used the difference of the average nuclear diameter across a number of randomly selected cells before and after expansion as a surrogate for expansion factor determination (Liffner et al., 2024) (Figure 1D and E).

**Figure 1:**
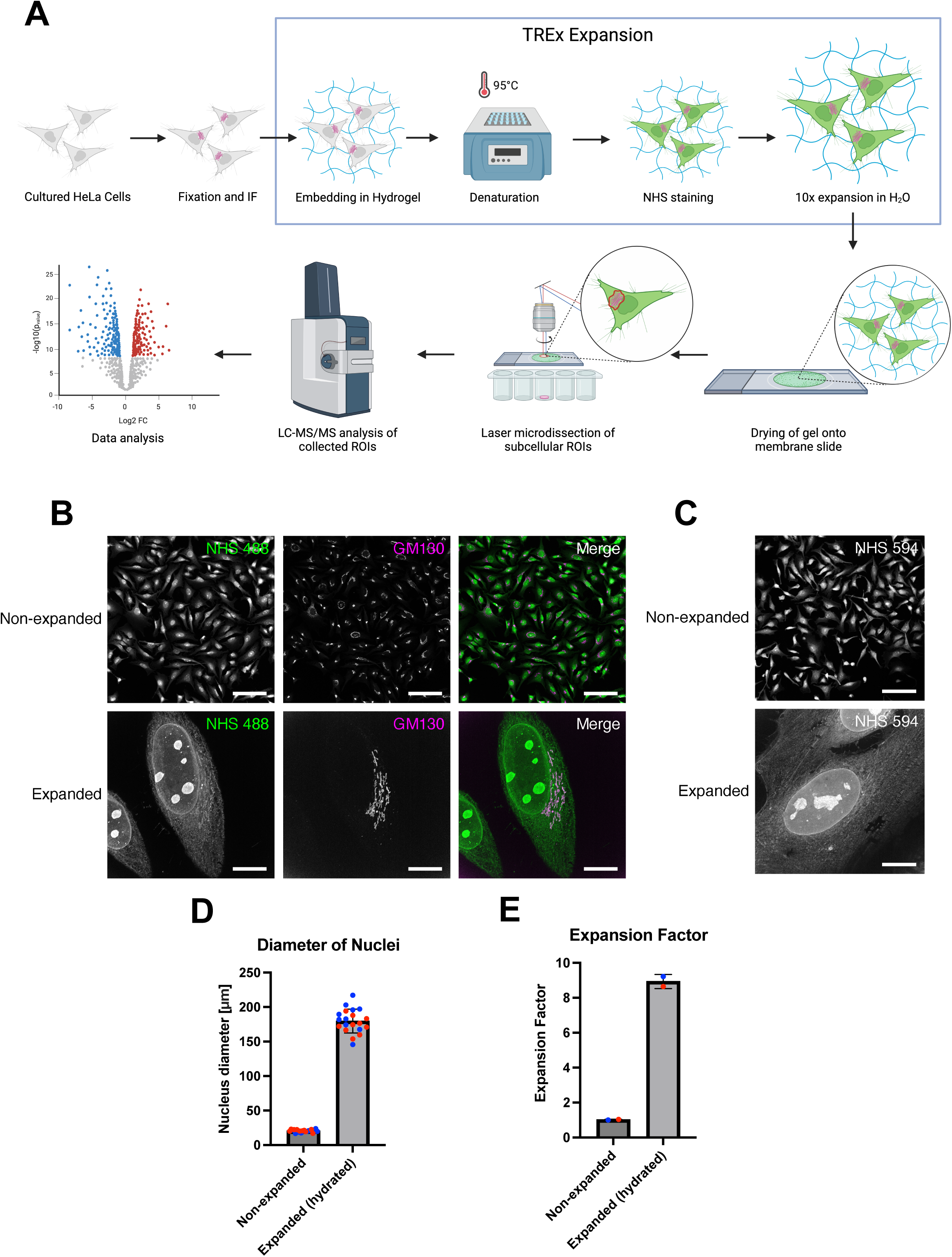
Development of the SPEx protocol. **(A)** SPEx workflow. Cultured HeLa cells are fixed, and immunofluorescence is used to label the subcellular compartment of interest with fluorescent antibodies. Next, the cells are embedded in a hydrogel. To allow for later expansion, proteins are denatured overnight at 95 °C. NHS staining is performed to non-specifically label proteins before expansion of the gel in H₂O. Expanded gels are dried onto PPS membrane slides, and laser microdissection of the areas of interest is performed. LC–MS of the collected areas and subsequent data analysis are used to identify proteins enriched in specific subcellular compartments. Created with BioRender.com. **(B)** Cells stained for total protein content with NHS 488 and immunostained with a GM130 antibody, with and without expansion using a modified TREx protocol. Scale bar: 100 µm. **(C)** Cells stained for total protein content with NHS 594, with and without expansion using a modified TREx protocol. Scale bar: 100 µm. **(D)** Measured nuclear diameters for non-expanded and expanded (hydrated) cells. Data are plotted as mean ± SD (non-expanded: 20.53 ± 1.94 µm; expanded (hydrated): 179.70 ± 17.26 µm) (n = 10 cells; N = 2 independent experiments indicated by different colors). **(E)** Average expansion factor calculated based on the measurements in (D). Data are plotted as mean ± SD (expanded (hydrated): 8.934 ± 0.40) (N = 2 independent experiments indicated by different colors).

#### Laser Microdissection

The aqueous properties of the gels were incompatible with laser cutting, due to laser diffraction and strong surface tension. To be able to collect organelles by laser microdissection, we completely dried the expanded gels on the polyphenylene sulfide (PPS) membrane slides. This procedure conserved the fluorescence signal from both, NHS dyes as well as secondary antibodies, which is used for LMD (Figure 2A and B). Conveniently, the drying of the gels did not lead to severe shrinkage in length and width (x-y direction), but only in height (z-direction) (Figure 2C, D and Figure S1B). Thus, we routinely retained an expansion of about 7x in x-y direction. The dried gels allowed efficient cutting at subcellular resolution using a Leica LMD7 system (Figure 2E, Supplementary video 1). We concentrated our efforts on three different organelles: the nucleus, the membraneless nucleolus, which were both nicely stained with the NHS ester, and the Golgi apparatus, visualized by anti-GM130. A summary of the number of cells and organelles, and which objective magnification was used to cut cells and organelles are outlined in Table S3. We did not only isolate these organelles from individual cells but also excised the exact same cells from which the nucleus/Golgi apparatus were removed (CellsMinusNucleus/CellsMinusGolgi) (Figure 2E). Similarly, after sampling the nucleoli, we also cut the nuclei from which the nucleoli were taken (NucleiMinusNucleoli) (Figure 2E). This was only possible because laser dissection allowed for repeated sampling from the same cell, without compromising sample quality.

**Figure 2:**
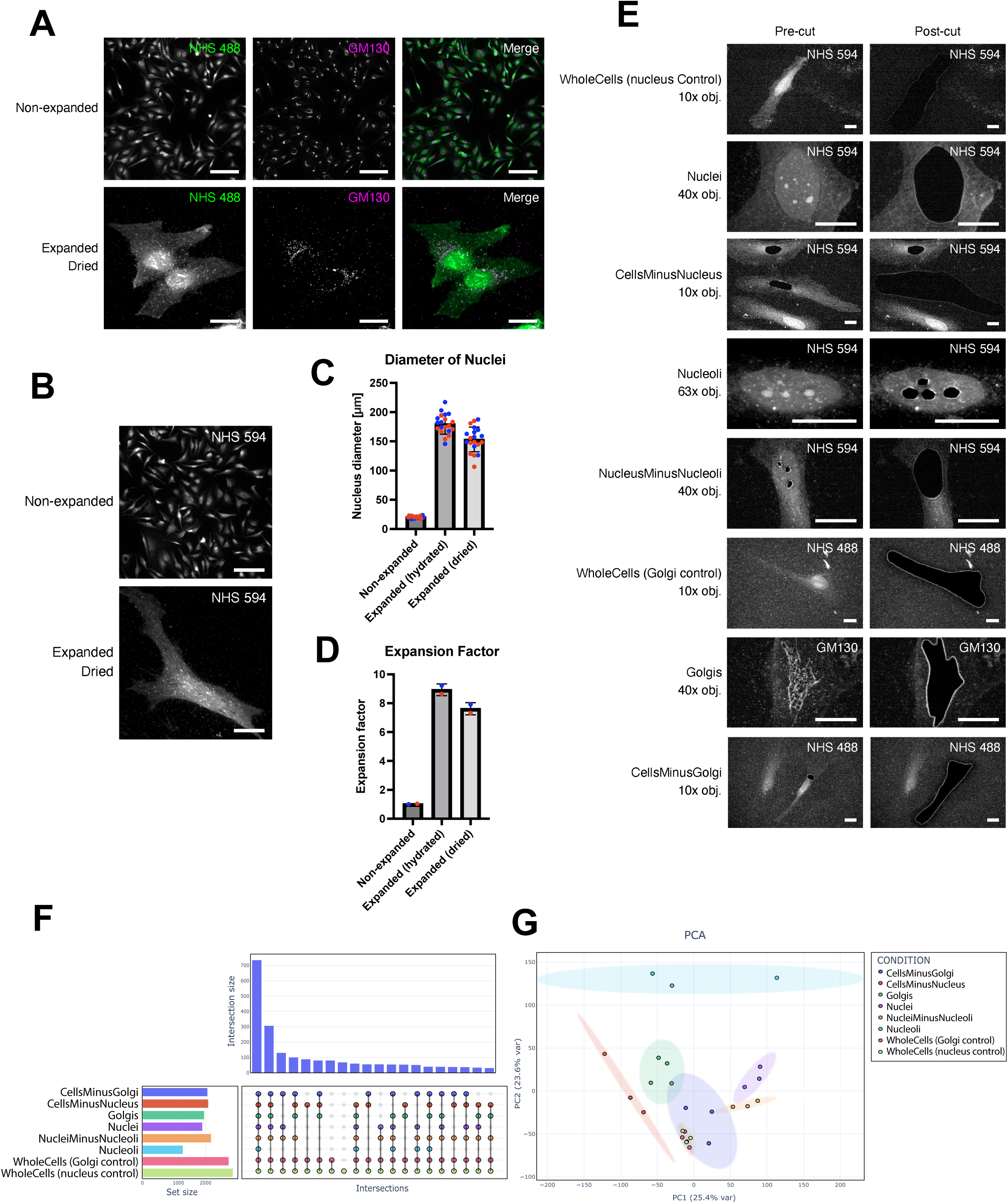
Optimization of the SPEx protocol for laser microdissection. **(A)** Cells stained for total protein with NHS 488 and immunostained for GM130, with and without expansion using a modified TREx protocol. Expanded samples were dried onto PPS membrane slides prior to imaging. Scale bar, 100 µm. **(B)** Cells stained for total protein with NHS 594, with and without expansion using a modified TREx protocol. Expanded samples were dried onto PPS membrane slides prior to imaging. Scale bar, 100 µm. **(C)** Nuclear diameters in non-expanded, expanded (hydrated and expanded (dried) cells. Data are mean ± s.d. (non-expanded: 20.53 ± 1.94 µm; expanded (hydrated): 179.70 ± 17.26 µm; expanded (dried): 153.30 ± 21.16 µm; n = 10 cells, N = 2 independent experiments indicated by different colors). **(D)** Expansion factor calculated from measurements in (C). Data are mean ± s.d. (expanded (hydrated): 8.934 ± 0.40; expanded (dried): 7.61 ± 0.42; N = 2 independent experiments indicated by different colors). **(E)** Pre- and post-cut images of all isolated sample types. Dissected cells were stained, expanded and dried according to the SPEx protocol. Images are not to scale (magnification of the objective used for dissection is indicated). Scale bar, 100 µm. **(F)** UpSet plot showing shared and unique proteins identified across the different regions of interest (ROIs) shown in (E). The horizontal bar chart on the left (Set Size) indicates the total number of proteins identified in each compartment. The vertical bar chart on the top (Intersection Size) displays the number of proteins specific to each unique intersection or combination of sets, represented by the matrix of black dots and connecting lines in the lower panel. Single filled dots represent set-specific (exclusive) proteins, while vertically aligned connected dots represent overlapping intersections between multiple groups. The degree of intersection size is sorted by frequency. Total number of analyzed proteins across all sets is N = 3021 (n = 3 or 4 biological replicates per group). **(G)** Principal component analysis (PCA) of proteomics samples from the same ROIs shown in (E). The scatter plot shows the first two principal components, PC1 and PC2, which account for 25.4% and 23.6% of the total variance, respectively. Each data point represents an individual biological sample (n = 3 or 4 per group). Colors and shapes indicate the distinct cell compartment groups. Ellipses represent the 95% confidence intervals for each group cluster.

#### LC-MS analysis

The collected samples were then prepared for proteomics, using an optimized protocol for low-input samples based on previous spatial proteomics workflows and latest generation of highly sensitive time-of-flight (ToF) LC-MS instrumentation using DIA-PASEF data acquisition (Makhmut et al., 2026; Rosenberger et al., 2025). Notably, we detected no artificial chemical lysine modifications from NHS-labeling or anchoring. In addition, we also analyzed the ratio of C-terminal lysine vs. arginine peptides and did not detect any bias (Figure S1C). This indicates that, if existent, such modifications are sparse with negligible proteome coverage impact. We consistently detected between 1,200 and 2,800 proteins per sample (Figure 2F and Tables S4 and S5). As expected, the number of identified proteins correlated with organelle size, where whole cells had the highest and nucleoli the lowest number of proteins. Moreover, principal component analysis (PCA) revealed very good clustering of biological replicates and, with the exception of whole cell controls, distinct segregation of the different sample types (Figure 2G). Of note, the whole cell samples from the nuclei and Golgi experiments cluster nicely together, confirming robust proteome quantification of our SPEx workflow. Taken together, SPEx allows the expansion of cells, laser dissection of membrane-bounded and membraneless organelles and the detection of protein constituents in these organelles.

### SPEx is suitable for the identification of nuclear proteins

To test the specificity and sensitivity of the SPEx method, we first applied it to the analysis of nuclear proteins from HeLa cells. The NHS ester allows the visualization of the entire cell and nucleus, and, hence, we cut out 25 nuclei from expanded cells and subjected them to LC-MS analysis. In addition, we analyzed the proteome of either 10 cells or 12 cells from which the nuclei had been dissected (CellsMinusNuclei). These samples were subjected to LC-MS analysis. First, we determined the percentage of the protein mass in the nuclei sample, which is contributed by proteins annotated to be located to the nucleus according to GO. We found that 75.5% of the total protein mass is contributed by nuclear proteins indicating good purity (Figure 3A). Next, we compared either whole cells or CellsMinusNuclei to the nuclei sample. As expected, we observed much stronger abundance differences, and therefore a higher number of significantly enriched nuclear proteins, when comparing nuclei to CellsMinusNuclei rather than to WholeCells (Figure 3B and S1D). Moreover, the comparison of the proteome of the nuclei to the one of the remaining cells without nucleus minimizes cellular proteome variability. Using this comparison, we quantified around 2,200 proteins from only 25 nuclei (Figure 2F and Table S5), with 522 proteins being significantly enriched in the nuclear samples. To evaluate the specificity of our approach, we checked the proportion of nuclear annotated proteins according to GO (Table S6) in the enriched protein fraction. We found 499 of the 522 proteins to be annotated nuclear, corresponding to a very high specificity of more than 95% (Figure 3C). Importantly, SPEx identified 23 proteins with strong nuclear enrichment that had not been annotated to be nuclear previously. Those proteins included SPOUT1 and C7orf50. Using immunofluorescence on non-expanded HeLa cells, we confirmed the nuclear localization of C7orf50 (Figure 3D) and SPOUT1 (Figure 4F), validating the high spatial specificity achieved with our approach. In line with our findings, recently C7orf50 has been identified as an oncogene with an N-terminal nuclear localization signal (Sun et al., 2026). Mechanistically, it has been proposed to shuttle between the nucleoplasm and the nucleolus, regulating autophagy and ribosome biogenesis downstream of mTOR (Sun et al., 2026). As an additional specificity test, we performed a functional enrichment analysis, which confirmed all top significantly enriched categories to be nuclear (Figure 3E and Table S7).

**Figure 3:**
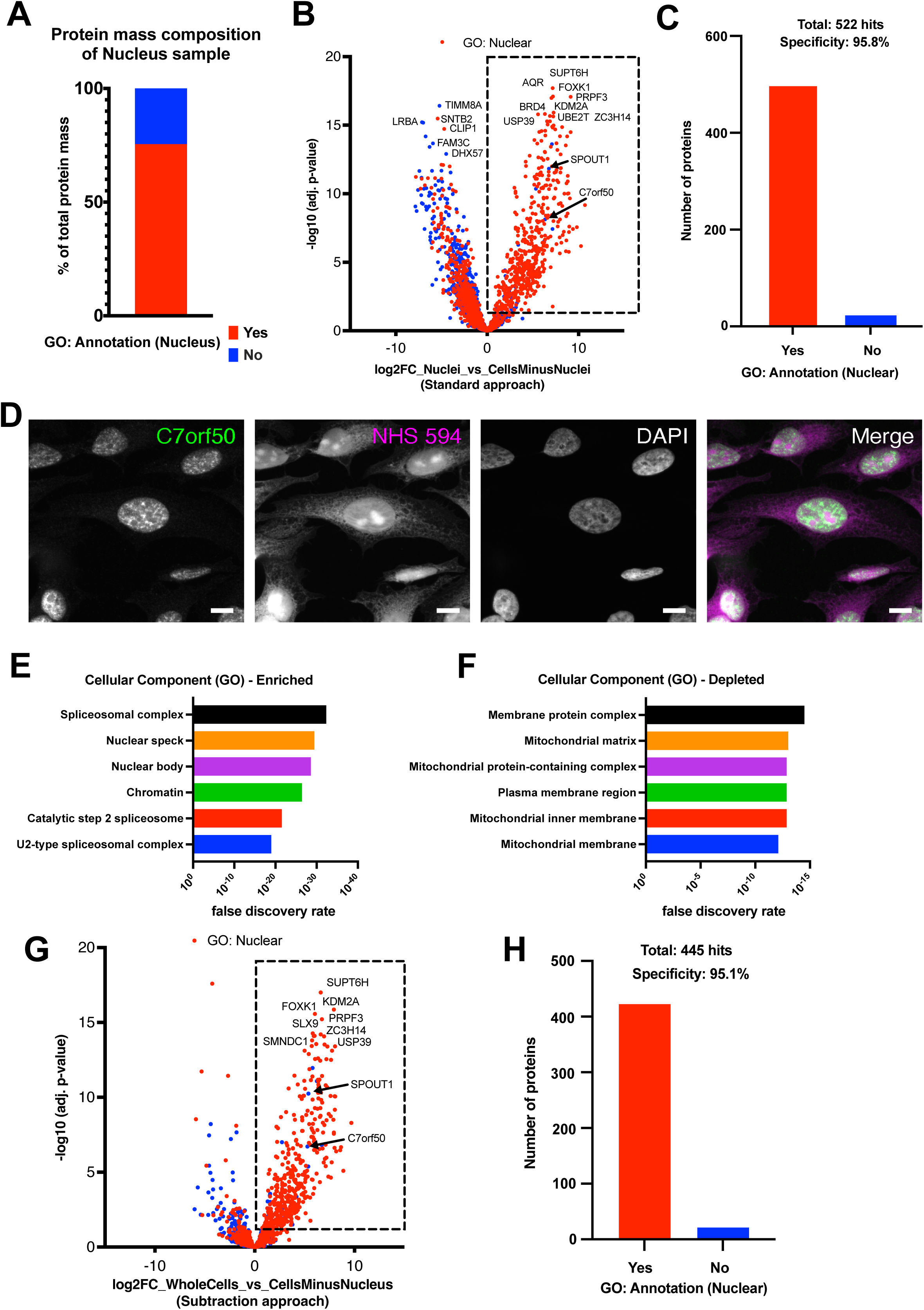
Application of SPEx to the nucleus. **(A)** Protein mass composition of nucleus samples. Red bar indicates the percentage of protein mass contributed by proteins annotated as nuclear according to Gene Ontology (GO) to the total protein mass based on intensity-Based Absolute Quantification (iBAQ) values. **(B)** Volcano plot of pairwise proteomic comparison between Nuclei and CellsMinusNuclei samples. The dotted square indicates proteins significantly enriched in the nucleus (adjusted *P* < 0.05, log₂FC > 0). Red dots represent proteins annotated as nuclear according to Gene Ontology (GO). FC, fold change. **(C)** Number of proteins significantly enriched in the nucleus in (B, dotted square) that are annotated as nuclear (Yes) or not nuclear (No) according to GO. **(D)** HeLa cells immunostained for C7orf50 and stained with DAPI and NHS 594. DAPI is omitted from the merged image for clarity. Scale bar, 10 µm. **(E)** Functional enrichment analysis of all proteins shown in (B) for cellular components using Gene Ontology (GO) via STRING (string-db.org). The six most significantly enriched terms (ranked by FDR) are shown. All significant terms (FDR < 1%) are listed in Table S7. **(F)** Same analysis as in (E), showing the six most significantly depleted GO terms. **(G)** Volcano plot of pairwise proteomic comparison between WholeCells and CellsMinusNucleus samples. The dotted square indicates proteins significantly enriched in WholeCells (adjusted *P* < 0.05, log₂FC > 0). Red dots represent proteins annotated as nuclear according to GO. **(H)** Same analysis as in (C) for proteins significantly enriched in WholeCells (G, dotted square).

**Figure 4:**
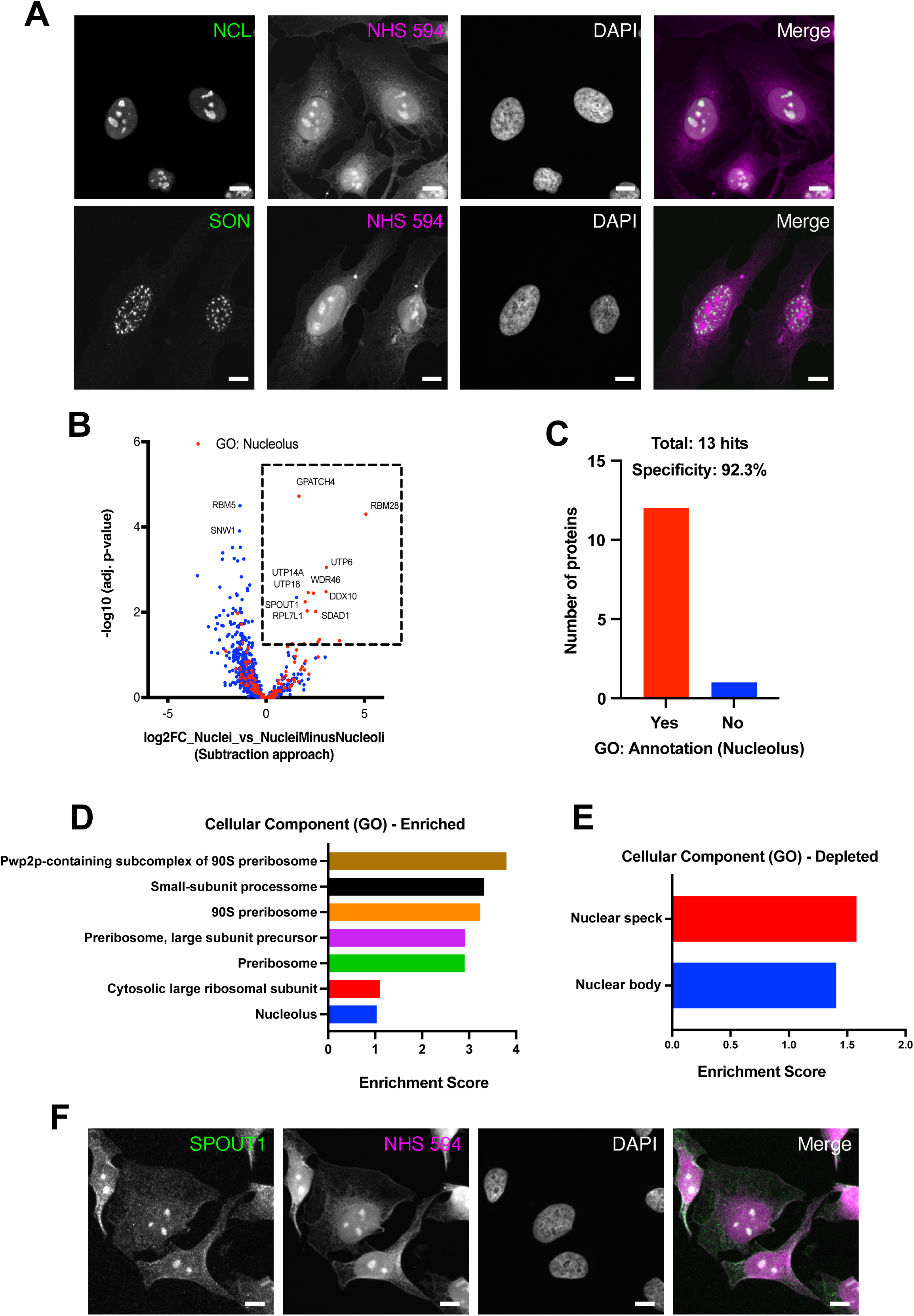
Application of SPEx to nucleoli. **(A)** HeLa cells immunostained for NCL (nucleolar marker) or SON (nuclear speckle marker) and stained with DAPI and NHS 594. DAPI is omitted from the merged image for clarity. Scale bar, 10 µm. **(B)** Volcano plot of pairwise proteomic comparison between Nuclei and NucleiMinusNucleoli samples. The dotted square indicates proteins significantly enriched in nuclei (adjusted *P* < 0.05, log₂FC > 0). Red dots represent proteins annotated to localize to the nucleolus according to Gene Ontology (GO). **(C)** Number of proteins significantly enriched in (B) that are annotated as nucleolar (Yes) or not (No) according to GO. **(D)** Functional enrichment analysis of all quantified proteins shown in (B) for cellular components using Gene Ontology (GO) via STRING (string-db.org). The seven most enriched significant terms (ranked by enrichment score) are shown. All significant terms (FDR < 1%) are listed in Table S11. **(E)** Same analysis as in (D), showing the two significantly depleted GO terms. **(F)** HeLa cells immunostained for SPOUT1 and stained with DAPI and NHS 594. DAPI is omitted from the merged image for clarity. Scale bar, 10 µm.

### Identification of proteins that are absent from the nucleus

Interestingly, functional enrichment analysis also identified significantly depleted cellular compartments, such as the plasma membrane and mitochondria (Figure 3F). Since cells without nuclei were used as a control for our analysis, SPEx can apparently not only be employed to localize proteins that are enriched in the nucleus but also detect those that are specifically depleted and hence absent from the nuclear fraction. This effect is already evident in the volcano plot, as proteins that are exclusively present in the cytoplasm (DHX57), peripherally associated with non-nuclear organelles (Golgi: LRBA and FAM3C) or constituents of those organelles (mitochondria: TIMM8A) are among the most significantly depleted proteins (Figure 3B).

Many proteins annotated to be nuclear by GO have either also functions in the cytoplasm and/or partition between the nucleus and other cellular locations. The quantitative capabilities of SPEx allowed to define the primary location of a protein. Even if a protein is annotated to be nuclear by GO, but depleted in the nuclei sample, this protein is most likely in higher amount outside than inside the nucleus. For example, CLIP1 is annotated to be nuclear using GO. However, in our analysis, CLIP1 was strongly depleted from the nuclei (Figure 3B). In agreement with our findings, CLIP1 (CLIP-170) is a microtubule binding protein in the cytoplasm (Perez et al., 1999), also annotated as such in uniprot, proteinatlas and opencell (Cho et al., 2022; Nirschl et al., 2016; “The Human Protein Atlas,” n.d.; The UniProt Consortium, 2025; Thul et al., 2017). Likewise, SNTB2 was annotated to be nuclear by GO. In contrast, it is annotated to be in the cytoplasm according to UniProt and has a confirmed localization in the cytoplasm/Golgi in the Human Protein Atlas. Moreover, SNTB2 has been localized to the cytoplasm in adipocytes (Krautbauer et al., 2019). Thus, we confidently detect proteins that are strongly depleted from the nucleus. To take this analysis a step further, we wanted to assess the specificity of detection of non-nuclear proteins in the depleted fraction. Therefore, we determined the proportion of proteins annotated to be mitochondrial and absent from the list of nuclear annotated proteins. Overall, 164 proteins annotated ‘mitochondrial’ were found in our dataset and all but one of them were reduced in abundance in our nuclear sample (Figure S1E). Only SPOUT1 was found to be significantly enriched in the nuclear fraction, but this protein is wrongly annotated as it is actually not a mitochondrial protein but instead present in the nucleus (Figure 4F). Taken together, SPEx cannot only be used to quantitatively identify proteins specifically enriched in an organelle, but also proteins, which concentrations are reduced in an organelle. The quantitative aspect of our approach makes it highly suitable for the analysis of changes in protein localization under different conditions.

### Determining the nuclear proteome via a subtraction approach

Encouraged by these results and as another test of the quantitative aspect of our method, we asked whether we could determine the proteome by subtraction (Figure S1F). Specifically, we wondered, if we compare the proteome of entire cells (WholeCells) to the one of cells from which we cut out the nucleus (CellsMinusNucleus), would we still be able to retrieve the nuclear proteome? We thought this might be feasible because the samples were well-separated in the PCA (Figure 2G). To our delight, this approach likewise showed a high enrichment of nuclear proteins (Figure 3G and H), achieving almost the same high specificity (95%) and coverage (423 nuclear proteins) compared to using the standard approach on the nuclei samples (Figure 3B and 3C). Moreover, the high overlap of the significantly enriched proteins confirmed the high specificity achieved with either comparison. In fact, the about 400 proteins, which were common between the two analyses (Figure S1G) showed an extraordinary nuclear specificity of 98% (Figure S1H). Thus, the subtraction method can also be used to reliably identify the proteome of a compartment, without analyzing the compartment itself. Moreover, combining the two quantitative analyses can be used to further boost the specificity of the SPEx method.

### SPEx compared to other spatial proteomics methods applied to the nucleus

To further assess the specificity and performance of our method, we next compared it to previously published large subcellular datasets, based on either subcellular fractionation followed by proteome profiling (Geladaki et al., 2019; Orre et al., 2019) or native immunoprecipitation proteomics of organelles (Hein et al., 2025) (Tables S8, S9 and S10). While the other methods could overall identify more proteins in their nuclear fractions (Figure S1I), SPEx achieved the highest specificity of all datasets (Figure S1J).

Taken together, SPEx is a new method to identify reliably and with high specificity nuclear proteins from low input samples. SPEx serves also as a discovery tool as we identified unannotated nuclear proteins. Moreover, our method is quantitative and efficiently determined proteins that were depleted or absent from nuclei.

### SPEx identifies the nucleolar proteome

To test whether SPEx also allows the determination of the proteome of much smaller structures within the cell, we aimed to investigate the proteome of nucleoli. Targeting nucleoli bears another challenge in that nucleoli are membraneless organelles, and many components are constantly exchanging between the nucleoplasm and the nucleoli, unlike when targeting a membrane-bounded organelle. Furthermore, the biochemical purification of membraneless organelles is challenging, and hence, the composition of the nucleolus is mostly based on imaging data or on GO annotation trained algorithm (Hein et al., 2025 Cell).

First, we aimed to confirm that the NHS ester-stained structures in the nucleus are indeed nucleoli. Therefore, we stained cells for nucleolin (NCL), an established nucleolar marker, or SON, which marks nuclear speckles (Hondele et al., 2019; Su et al., 2013). Indeed, NHS ester marks specifically nucleoli and not nuclear speckles (Figure 4A). We therefore cut out 100 nucleoli and subsequently the corresponding nuclei without nucleoli (NucleiMinusNucleoli) and compared their proteomes (Figure S2A). We detect about 300 proteins, of which according to GO only 11% were annotated to be nucleolar localized (Figure S2B). This was also reflected in the mass contribution of proteins which are annotated as nucleolar according to GO, which was only 3.9% (Figure S2C). One reason could be that most nucleolar protein shuttle between the nucleoli and the nucleoplasm, and therefore their abundance could be rather similar in the nucleoplasm and the nucleoli, while their concentration could be still higher in the nucleoli. Moreover, the volume of the nucleus is much bigger than the combined volume of the nucleoli, which we excised, making quantitative comparison more challenging. To circumvent these problems, we applied the subtraction method to detect which proteins are reduced in the NucleiMinusNucleoli samples compared to the Nuclei samples. Using this method, we increased the specificity from 10.8% to 92.3%, at the expense of number of specific hits (Figure 4B). Still, from the 13 significantly enriched proteins all but one, SPOUT1, were annotated by GO to be nucleolar (Figure 4C). We have already identified SPOUT1 as a novel nuclear protein above (Figure 3B). Our nucleoli analysis suggested that SPOUT1 is actually a nucleolar protein that we could confirm by immunofluorescence (Figure 4F). This is in line with a study showing that the SPOUT1 homolog in drosophila (Ptch) is a methyltransferase that methylates the 28S rRNA in the Drosophila nucleolus and is essential for oogenesis (Chen et al., 2024). Moreover, functional enrichment analysis showed that proteins associated with the 90S pre-ribosome, which are specifically localized in the nucleolus, were highly enriched (Figure 4D and Table S11). Thus, despite this high stringency, we were still able to identify a novel component of the nucleolus. It is worthwhile noting that while the subtraction method strongly reduced the number of potential false positive, the most significant enriched proteins were also found by the standard method in which we compared Nucleoli to NucleiMinusNucleoli (Figure 4B and S2A), indicating that nucleoli proteins were also enriched in the excised nucleoli fraction. Moreover, the standard method might at times be more useful as a discovery tool. Nevertheless, we successfully identified proteins of the membraneless nucleolus using expanded *in situ* laser-dissected samples.

### SPEx identifies nuclear speckle proteins absent from nucleoli

The nucleolus is not the only membraneless organelle in the nucleus. Like nucleoli, those other organelles, such as nuclear speckles, have a high local concentration of their protein components, but these proteins are likewise present in the nucleoplasm, and both pools are constantly exchanging. However, exchange between constituents of nucleoli with constituents of other nuclear membraneless organelles is expected to be limited. As a further test for specificity, we asked whether we could detect components of the other nuclear membraneless organelles in the fraction that is depleted or absent from nucleoli. In both, the standard and the subtraction method, RBM5 and SNW1 were very significantly depleted from the nucleoli fraction (Figure 4B and S2A). RBM5 is a component of the spliceosome A complex and SNW1 a component of the minor spliceosome. The spliceosome has been shown to be located in nuclear speckles (Bhat et al., 2024) and, according to protein atlas (Figures S2D and S2E), RBM5 and SNW1 are indeed located in nuclear speckles and absent from the nucleolus. Moreover, functional enrichment analysis revealed that nuclear speckle and nuclear body proteins were significantly depleted from our nucleolar fraction (Figure 4E and Table S11). Therefore, also for the nucleolus, we were able to determine proteins that were specifically depleted/absent from nucleoli in an environment, which is guided by protein diffusion.

### SPEx compared to other spatial proteomics methods applied to the nucleolus

To date, only few subcellular proteome studies assigned nucleolar locations to proteins, because membraneless organelles are notoriously difficult to purify (Orre et al., 2019) (Table S8). By contrast, SPEx allows isolation and collection of organelles directly from cells and is therefore ideally suited for the enrichment and analysis of membraneless organelles. This is nicely demonstrated by the higher specificity for nucleolar proteins obtained by SPEx (>90%) compared to classical density gradient centrifugation with only around 50% (Orre et al., 2019) (Figure S2F and S2G).

Taken together, SPEx not only allowed us to detect nucleolar proteins with very high specificity, we also identified an unannotated nucleolar protein. Moreover, compared to other available methods, SPEx demonstrated a higher specificity assignment of proteins for a membraneless organelle, like the nucleolus.

### Golgi proteins are identified with high specificity using SPEx

Finally, we used SPEx to determine the proteome of the Golgi. The Golgi is a central sorting station for proteins and lipids and which is responsible for post-translational modifications such as glycosylation, sphingolipid synthesis and sulfation. It is a highly dynamic structure, and transport through the Golgi and the dynamics of Golgi cisternae are still a matter of debate. To address these issues, it would be desirable to have a method, which could capture the Golgi in different states. Recently developed immunoprecipitation methods have tackled this challenge (Fasimoye et al., 2023), however, because of a few time-consuming purification steps, important transient molecular interactions might still be lost. SPEx, on the other hand, allows direct isolation of organelles from fixed cells with preserved molecular composition and should therefore be well suited for the analysis of this dynamic organelle.

To test this exciting possibility, we excised 35 Golgis from expanded and dried samples and performed LC-MS. First, we determined which percentage of the total protein mass in these samples is contributed by proteins GO annotated with Golgi or extracellular localization. The later class most likely represents cargo traveling through the Golgi. We found that 84.7% of the total protein mass is contributed by proteins belonging to one or both of those classes, indicating very good purity of the sample (Figure S3A). Next, we compared these Golgi samples to the corresponding cells without Golgi (CellsMinusGolgi) (Figure 5A). We identified 356 significantly enriched proteins in the Golgi samples, 288 (80.9%) were GO annotated with Golgi or extracellular localization (Figure 5B). Among the enriched proteins were many proteins related to glycosylation such as GALNT1,2 and 7 (Figure 5A). We also detected several Golgins (GOLGA2-4); GOLGA2 (GM130) was our Golgi marker. Moreover, numerous trafficking-related Golgi proteins such as the SNARE receptor GOSR1, the COG tethering proteins and the coatomer complex, which is the coat of COPI vesicles, were highly enriched in the Golgi fraction. The method was sensitive enough to also detect the Golgi Rab, Rab6A (Figure 5A). In contrast, endosomal proteins such as SNX2, CHMP4A and B or EHD2 were depleted as well as the components of the adaptor complex for clathrin-mediated endocytosis such as AP2A2 or the plasma membrane exocyst tether EXOC8 (Figure 5A). Interestingly, we also observed depletion of the ARP2/3 complex from the Golgi fraction. Notably, the comparison of WholeCells to CellsMinusGolgi revealed also Golgi-resident proteins (Figure S3B), but with a somewhat lower specificity (Figure S3C). The trend of the of the analysis is similar to what we observed for the analysis of the nucleus (WholeCells vs CellsMinusNucleus). Functional enrichment analysis confirmed the high specificity of Golgi proteins in our SPEx results. The most enriched cellular components were all associated with the Golgi (Figure 5C and Table S12), whereas the most depleted protein groups were all assigned to the nucleus (Figure 5D). This result may not be very surprising, because the nucleus has the highest protein content and there is essentially no overlap with Golgi proteins.

**Figure 5:**
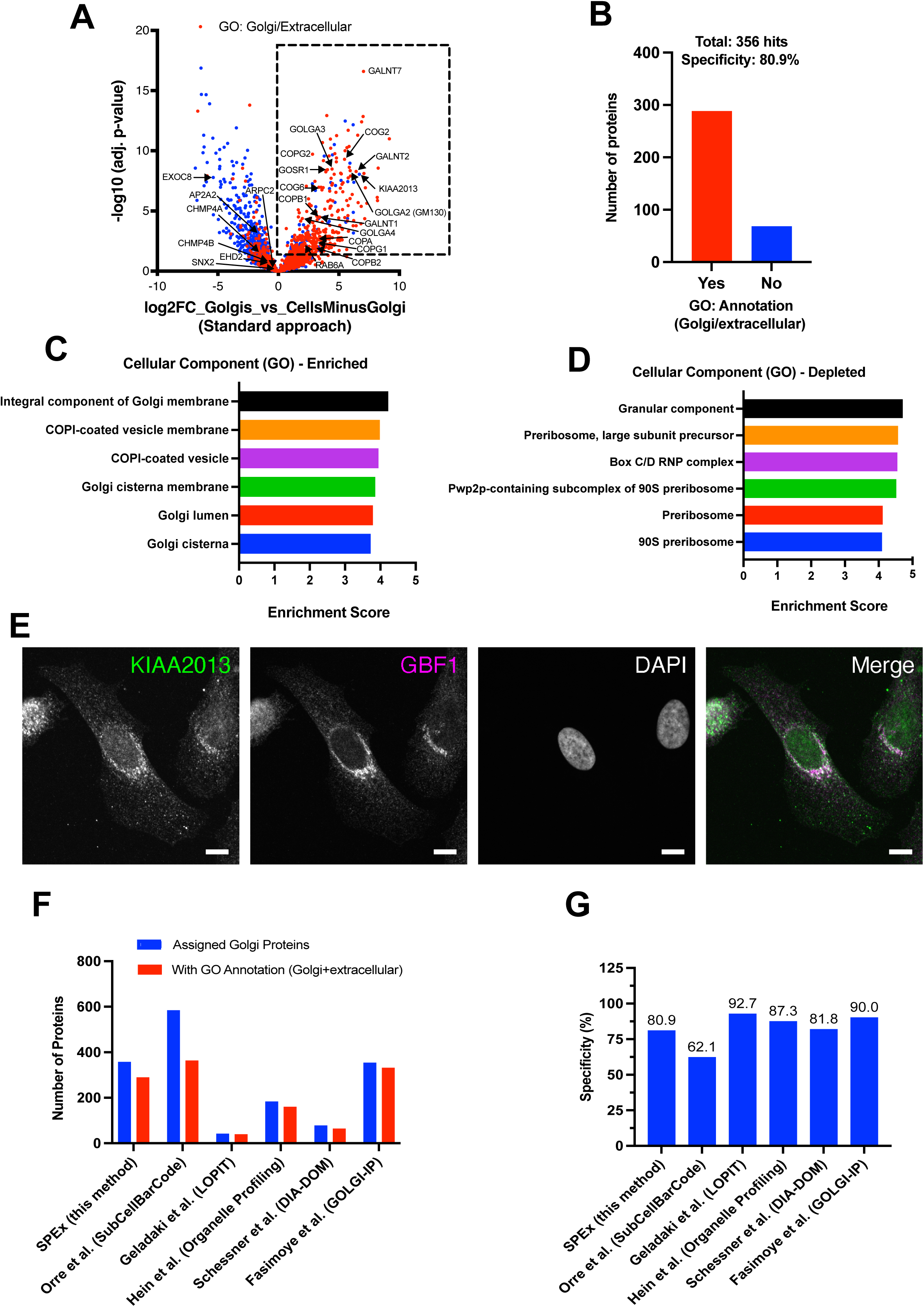
Application of SPEx to the Golgi. **(A)** Volcano plot of pairwise proteomic comparison between Golgis and CellsMinusGolgi. The dotted square indicates proteins significantly enriched (adjusted *P* < 0.05, log₂FC > 0) in the Golgi. Red dots represent proteins annotated to localize to the Golgi and/or to be extracellular according to GO. **(B)** Number of proteins significantly enriched in the Golgi in (A) that are annotated as Golgi and/or extracellular (Yes) or not (No) according to GO. **(C)** Functional enrichment analysis of all quantified proteins shown in (A) for cellular components using Gene Ontology (GO) via STRING (string-db.org). The six most enriched terms (ranked by enrichment score, FDR < 1%) are shown. All significant terms (FDR < 1%) are listed in Supplementary Table S12. **(D)** Same analysis as in (C), showing the six most depleted GO terms. **(E)** HeLa cells immunostained with antibodies for KIAA2013 and GBF1 (Golgi marker) and stained with DAPI. DAPI is not shown in the merged image for clarity. Scale bar, 10 µm. **(F)** Bar plot comparing the number of proteins assigned to the Golgi (blue) in our SPEx dataset (this study, A) with other published spatial proteomics datasets (Fasimoye et al., 2023; Geladaki et al., 2019; Hein et al., 2025; Orre et al., 2019; Schessner et al., 2023). Proteins overlapping with GO Golgi/extracellular annotations are shown in red. **(G)** Specificity of GO Golgi/extracellular annotation in the different proteomics datasets, determined from (F).

Our analysis also identified proteins that were not annotated previously to be Golgi localized. We selected again one candidate, KIAA2013, for confirmation (Figure 5A) and determined the subcellular localization using immunofluorescence (Figure 5E). Indeed, KIAA2013 was Golgi localized and, thus, again we were able to assign a new location to a protein. KIAA2013 remains very poorly characterized but based on sequence analysis the protein contains a short C-terminal cytoplasmic tail, a single transmembrane domain and a large, potentially extracellular, N-terminal domain (The UniProt Consortium, 2025). While our SPEx and immunofluorescence data suggests that the bulk of KIAA2013 is retained at the Golgi, the additional cytoplasmic puncta raise the possibility that a fraction of KIAA2013 is trafficked beyond the Golgi, potentially to the plasma membrane or to endosomal compartments.

### SPEx compared to other spatial proteomics methods

Compared to previously published large scale datasets (Fasimoye et al., 2023; Geladaki et al., 2019; Hein et al., 2025; Orre et al., 2019; Schessner et al., 2023) (Tables S8, S9, S10, S13 and S14), SPEx achieved comparable results with a high proteome coverage (Figure 5F) and similar specificity (Figure 5G). Finally, we also compared SPEx to previously published methods combining ExM and spatial proteomics (Dong et al., 2024a; Li et al., 2022) (Table S15). SPEx outperforms previously available methods with respect to spatial resolution. Thus, SPEx is the first method that allows LMD and LC-MS of structures significantly smaller than the nucleus, while maintaining good proteome coverage.

## Discussion

Here, we report the development of SPEx as a method for *in situ* subcellular spatial proteomics at a single micrometer resolution. To achieve this, we combined expansion microscopy with laser dissection and LC-MS/MS. SPEx requires low sample input, is low cost and adaptable to membrane-bounded and membraneless organelles. Depending on the size of the organelles, SPEx can be readily used for subcompartments of organelles.

### Development of SPEx

To establish the protocol, we took advantage of the well-known TREx protocol, which allows up to 10x expansion without the need for any specialized equipment or procedures (Damstra et al., 2022). Drying the samples onto a PPS-membrane still maintained an about 7x linear expansion in x-y, resulting in almost 50x increase of surface area. This increase enlarges the subcellular regions to an extent that they can be easily excised by laser microdissection. Depending on the size of the structure of interest, SPEx is also compatible with lower expansion factors, which can be fine-tuned by adding different amounts of the crosslinker (Bis) to the gelation solution. For example, we reached 4-fold, 6-fold or 10-fold expansion by using 0.1%, 0.04% or 0.005% Bis, respectively. In this study, we focused on 10x expansion since it enables isolation of structures as small as nucleoli. In addition, we found that small differences in the amount of anchoring reagent (AcX) impacted the expansion factor of cells but not of the hydrogel. For this reason, we reduced anchoring from 0.1 mg/ml AcX to 0.05mg/ml AcX compared to (Damstra et al., 2022). This reduction showed no negative effect on proteome coverage in LC/MS but improved expansion factor. In case the expansion factor is not consistent between sample and gel, we suggest titrating AcX concentrations between 0.01 mg/ml and 0.1 mg/ml, to find optimal conditions. Moreover, the use of low amounts of glutaraldehyde (0.1%) during fixation is compatible with SPEx but might have a potential negative impact on the expansion factor due to increased crosslinking of the sample. Therefore, we would recommend the use of PFA only, whenever possible. We also found that secondary antibodies and dyes in the red-emitting range (e.g. 594 nm) are likely to be better in the preservation of fluorescence throughout expansion and drying of the sample. We, therefore, usually visualized the most relevant structure in the red-emitting range. Testing other secondary antibodies and labeling techniques should provide more additional color options, which might be useful for double labeling experiments. Such double labeling will facilitate the determination of the proteome of membrane subcompartments such as membrane contact sites. Finally, SPEx might be compatible with other protease free expansion workflows. However, we have not used other expansion methods, and compatibility would have to be assessed.

Another possible problem we considered is that lysine modifications during anchoring (or NHS labeling) might interfere with detection of the modified peptides during LC-MS and thus reduce coverage. We specifically searched for both modifications (NHS labeling and anchoring) in our LC-MS data but could not identify any such lysine modified peptides. Therefore, these modifications are either very sparse or not compatible with LC-MS analysis. To exclude the latter, we compared the ratio of peptides carrying a lysine at the last position to the peptides carrying an arginine at the last position. Since both amino acids are equally abundant in the human proteome and tryptic cleavage sites, lysine modifications will cause frequent miscleavage/prevent detection and we would expect a strong bias towards peptides carrying an arginine at the last position. However, we did not observe such a bias (Figure S1C) and obtained very similar Lys/Arg ratios compared to other fixed, but not expanded, human samples, indicating that our expansion protocol has no detectable impact on lysine residues.

### SPEx can be applied to a variety of subcellular targets

To demonstrate the general applicability of SPEx we chose a diverse set of targets in this study: the nucleus, the nucleolus and the Golgi. Those targets differ in size, shape, labeling method required for visualization (dyes and IF) and architecture (membrane-bound and membraneless organelles). Surprisingly only a couple of dozens of organelles are required as sample input for LC-MS/MS for SPEx. These inputs were sufficient to obtain good coverage and high specificity. Importantly, we detected not only predicted or previously known components of these well-studied organelles. We also identified novel proteins residing in the nucleus, the nucleolus and the Golgi with high specificity. Of note, membraneless organelles are inherently difficult to purify. SPEx overcomes this problem by eliminating the need of biochemical enrichment procedures. Moreover, SPEx compares favorably to other specialized enrichment procedures. We compared the coverage and specificity of SPEx to a number of currently available other spatial proteomics methods, based on either subcellular fractionation followed by proteome profiling (Geladaki et al., 2019; Orre et al., 2019; Schessner et al., 2023) or native immunoprecipitation proteomics of organelles (Fasimoye et al., 2023; Hein et al., 2025). When applied to the nucleus or nucleoli, SPEx showed either better coverage or better specificity than currently available methods. When we employed SPEx to the Golgi, we were able to obtain coverage and specificity comparable to currently available methods including a specific Golgi-IP (Fasimoye et al., 2023). Another great advantage of SPEx is that it does not rely on transfection of fusion proteins, changes in the media or other manipulations. Whatever method can be used to visualize the region or structure of interest is sufficient to perform SPEx. We used the same overall protocol for the sample preparation and the analysis of the nucleus, nucleoli and Golgi, indicating that not much, if any, tinkering must be done to perform a successful SPEx experiment. Sampling of different structures from the same preparation (e.g. whole cells, nucleus, cells without nucleus, nucleoli, nuclei without nucleoli) abolishes the need of separate sample preparations for different targets and controls. Compared to other methods utilizing ExM to perform spatial proteomics (Dong et al., 2024b; Li et al., 2022), SPEx exhibits a significant improvement in spatial resolution, thus allowing the analysis of a large number of subcellular compartments.

### Different analysis pipelines for SPEx data

Our data show that SPEx data can be analyzed efficiently in two different ways. The standard approach is to isolate structures of interest and compare their proteome to the one of cells or subcellular compartments from which the structures were removed (control). The other, useful option is the subtraction approach, which is an indirect measure, but still very powerful. In this case, the structure of interest is cut out. The analysis, however, is performed on the cell/organelle from which the structure of interest has been excised and compared to the cell/organelle in which the structure is still present. Both approaches yield comparable results in our test cases.

The standard approach is well-suited for most questions and approaches, and also for the discovery of transiently interacting or low abundant proteins. Even though we employed median normalization to our proteomics data abundances which controls for different sample amounts and eventually results in a comparison of protein concentration rather than pure protein abundances, it became clear that comparison of structures of very different sizes might be technically difficult and that in such cases, controls should be collected in a similar size range to the actually studied compartments. The subtraction method detects proteins that were lost from one sample, and is thereby highly specific, and gives less false positives. This approach might be particularly useful when venturing into the unknown and markers of a compartment or domain should be identified. Even though we obtain about 7x fold expansion of our dried samples, this magnification might not be sufficient to cut out and analyze tiny organellar subcompartments. In those cases, eliminating these structures from the sample might provide the specificity to identify key proteins of these structures.

### SPEx performs best in a comparative approach

We have great resolution in x-y direction (50x surface area increase), however no resolution in z. We were, therefore, initially worried about potential contamination and false positive detection of proteins. To our delight, it turned out that this was not a big problem. Proteins present on an organelle will still be enriched in the sample, while the level of the cytoplasmic or plasma membrane proteins will remain at the same levels in the sample and the control, and hence these proteins are not enriched in the organellar fraction. Even though, we may also cut out other organelles or pieces thereof, this will likewise happen in the control. For the Nucleus and the Golgi we also found that the vast majority of the protein mass in the sample is still contributed by proteins assigned to these respective compartments, indicating that even without comparing the samples to a control the data would represent an albeit less clean proteome of the organelle. For much smaller compartments like the nucleolus, however, this comparative approach is likely essential. This is illustrated by the fact that even though only a small part of the protein mass in our nucleoli samples was contributed by nucleolar proteins, comparison with a control allowed us to identify the same proteins as with the more stringent subtraction approach, however with lower specificity. Our data show that the lack of resolution in Z is a much lesser problem than initially anticipated.

### Future directions

We think that SPEx holds great potential to answer spatial proteomics questions that were hard to tackle until to date. A great strength of SPEx is that the experimentalist specifically selects the (sub) compartments for the analysis, which is in marked difference to proximity labeling techniques. SPEx could, thus, be used to quantitatively determine the composition of fragmented versus fused mitochondria or between tubular and sheet endoplasmic reticulum in the same cell. We foresee also the use to detect other organellar subcompartments. Moreover, compared to other spatial proteomics methods currently used, SPEx requires very little prior knowledge and no genetic manipulation of the studied system. Visualizing an organelle or compartment just by morphology would be sufficient for SPEx. Thus, even structures for which no antibodies are available could potentially be analyzed using SPEx. This, together with the fact that the TREx expansion method has been shown to be widely applicable to different cell types and tissue sections (Damstra et al., 2022), leads us to believe that SPEx could be a valuable tool to perform spatial proteomics on hard to transfect cells, tissues and non-model organisms were genetic manipulation and/or antibody staining remains highly challenging.

Another potential application of SPEx lies in studying cell to cell heterogeneity, because SPEx allows to not only perform spatial proteomics on specific organelles, but also allows researchers to select from which cells these organelles should be isolated. This would allow for comparative studies, where the proteome of specific organelles in morphologically different cells in the same dish (e.g. dividing vs non-dividing or responders vs non-responders in drug treatment) could be determined. This remains challenging with conventional methods, due to the generation of lysates from many cells prior to enrichment. Therefore, the analysis on the single cell level remains difficult with these methods. Importantly, SPEx does allow these types of analyses.

It is conceivable that the SPEx approach could be used beyond the context of spatial proteomics. In the future, this approach could potentially be coupled to downstream lipidomics allowing to study the membrane composition of different organelles/domains in the cell. Possibly, the same workflow could also be used to perform RNA/DNA sequencing to determine the RNA content of condensates or the DNA content of specific nuclear regions. We also expect to SPEx to work on tissue sections.

In summary, by combining ExM, LMD and low input LC-MS, SPEx offers a novel approach to subcellular spatial proteomics. The low sample requirements and flexible nature of the SPEx method make it a useful approach to tackle spatial proteomics questions that were previously hard to investigate and required the development of highly specialized methods.

## Material and methods

Key resource table.

Key resource table
| Reagent type or resource | Designation | Source or reference | Identifiers | Additional information |
| --- | --- | --- | --- | --- |
| Antibody | NCL (Mouse Monoclonal) | Abcam | ab136649 | Dilution (1:500) |
| Antibody | SON (Rabbit Polyclonal) | GeneTex | GTX123778 | Dilution (1:5000) |
| Antibody | GOLGA2/GM130 (Rabbit Polyclonal) | Proteintech | 11308-1-AP | Dilution (1:200) |
| Antibody | SPOUT1 (Rabbit Polyclonal) | Atlas Antibodies | HPA022990 | Dilution (1:50) |
| Antibody | KIAA2013 | Atlas antibodies | HPA060536 | Dilution (1:10) |
| Antibody | GBF1 (Mouse Monoclonal) | BD Biosciences | AB_399487 | Dilution (1:200) |
| Antibody | C7ORF50 | Atlas Antibodies | HPA052281 | Dilution (1:65) |
| Antibody | Rabbit IgG Alexa Fluor 594 (Goat Polyclonal) | ThermoFisher | A-11012 | Dilution (1:200) for ExM, Dilution (1:500) for IF |
| Antibody | Mouse IgG Alexa Fluor 594 (Goat Polyclonal) | ThermoFisher | A-11005 | Dilution (1:500) |
| Antibody | Rabbit IgG Alexa Fluor 488 (Goat Polyclonal) | ThermoFisher | A-11008 | Dilution (1:500) |
| Antibody | Mouse IgG Alexa<br>Fluor 488 (Goat<br>Polyclonal) | ThermoFisher | A-11001 | Dilution (1:500) |
| Cell line (homo<br>sapiens) | HeLa CCL2 | Gift from Dr.<br>Martin Spiess |  |  |

### Cell culture

HeLa cells were cultured at 37°C and 5% CO_2_ in growth media (DMEM high-glucose medium (Sigma-Aldrich, D57969) with 10% FCS (Biowest, S1810), 1% penicillin–streptomycin (Sigma-Aldrich, P4333), 1 mM sodium pyruvate (Sigma-Aldrich, S8636) and 2 mM L-glutamine (Gibco, 25030-024)). Cells were regularly tested for mycoplasma using a Mycoplasma PCR detection kit (Applied Biological Materials, G238).

### Fixation and immunofluorescence

50,000 HeLa Cells were seeded on sterile 13 mm glass coverslips (VWR, 631-0150) in a 24-well plate (Sarstedt, 83.3922). Cells were incubated at 37°C, 5% CO_2_ O/N, washed once in PBS (Gibco, 20012-019) and fixed in 3% PFA (Sigma, F8775-500ML) in PBS for 10 minutes. Cells were washed 3x in PBS and permeabilized with 0.1% Triton (Sigma, T8787-50ML) in PBS for 10 minutes. After permeabilization, cells were washed 3x in PBS and incubated in blocking solution (1% BSA (Sigma-Aldrich, A9647-100G) in PBS) for 30 minutes. Coverslips were transferred to a wet chamber and incubated for 1 hour in primary antibody (for dilutions see resource table) in blocking solution. Coverslips were washed 3x in PBS and incubated for 30 minutes in secondary antibody in blocking solution (for dilutions see resource table). Cells were washed 3x in PBS. Cells were incubated with 1 μg/ml DAPI (Sigma-Aldrich, D9542-10MG), and 0.2 μg/ml NHS ATTO 594 (Sigma-Aldrich, 08741-1MG-F) or NHS-ATTO 488 (Sigma, 41698-1MG-F), where applicable, in PBS for 1 hour at 37 °C. After washing 3x in PBS for 5 minutes, coverslips were mounted on Fluoromount-G (SouthernBiotech, 0100-01) and left to settle overnight in the dark. Unless mentioned otherwise, all steps were performed at room temperature (RT). Confocal images were acquired with Evident/Olympus Fluoview FV3000 laser scanning system equipped with solid-state lasers and a galvanometer scanner using an UPLSAPO 60×/1.30 objective with silicone oil (Evident/Olympus, Z81114) and FV31S-SW software.

### SPEx Workflow

#### Fixation and immunofluorescence for SPEx

Sterile 13 mm coverslips were prepared in a 24-well plate. Coverslips were coated by incubating them in coating solution (Fibronectin bovine plasma (Sigma-Aldrich, F1141-5MG) 1:200 in ddH_2_O) for 90 minutes at RT. The coating solution was aspirated, and the coverslip were air dried for 20 minutes and washed once in PBS. 20,000 HeLa cells were seeded in DMEM (low confluency (∼30-40%) is crucial for optimal expansion) and incubated at 37°C, 5% CO_2_ O/N.

Cells were washed once in PBS and fixed for 10 minutes at 37°C with pre-warmed fixation mix (4% PFA in PBS). Cells were washed 2x in PBS. If no IF was performed cells were permeabilized for 10 minutes in 0.1% Triton in PBS and subsequently washed 3x in PBS (for non-IF samples proceed to “TREx expansion”). For IF, cells were blocked for 1 hour in blocking solution (3% BSA, 0.1% Triton in PBS) with gentle rocking. Next, cells were incubated with primary antibody (for dilutions see resource table) in blocking solution for 3 hours with gentle rocking. Cells were washed 3x 5 minutes in PBS with gentle rocking and incubated with secondary antibody (for dilutions see resource table) in blocking buffer O/N at 4°C with gentle rocking. Cells were washed 3x 10 minutes in PBS with gentle rocking. Unless mentioned otherwise, all steps were performed at RT. Adapted from (Damstra et al., 2022).

#### TREx Expansion

Prior to the expansion, gelation chambers and gelation solution were prepared as follows:

• Gelation chamber: Two holes of 3-4 mm diameter (spaced apart by ∼3mm) were punched into a microscopy-slide-sized piece of parafilm (Parafilm, PM-996) using a tissue puncher. The parafilm was then placed on a glass microscopy slide (Epredia, AG00008032E01MNZ20) (paper side up) and adhered by repeatedly streaking over it with a second microscopy slide and removing the paper on the parafilm. Finally, streaking can be repeated without the paper attached, by putting a non-adhesive plastic foil over the parafilm and streaking over it again until the parafilm is perfectly adhered to the glass microscopy slide (Figure S3D, steps 1-4).

• Gelation solution: The base of the gelation solution (1.1 M sodium acrylate (Sigma, 408220-25G), 2 M acrylamide (Sigma, A4058-100ML), 0.005% Bis (Sigma, M1533-25ML), 1x PBS (Thermo Fischer, 70011044)) was mixed, aliquoted and stored at -20°C. Before gelation aliquots were thawed to RT (prevents incomplete sodium acrylate dissolution) and then placed on ice. Right before use 0.15% TEMED (Sigma, S7653-250G) and 0.15% APS (Sigma, A3678-25g) were added, the solution was mixed thoroughly and placed back on ice immediately.

Anchoring was performed by incubating cells for 1 hour in 0.05 mg/ml AcX (Invitrogen, A20770) in PBS. In case expansion of the gel and the sample are not equal, we suggest titrating AcX concentrations between 0.01 mg/ml and 0.1mg/ml for optimal results. Cells were washed once in PBS. The coverslip was rinsed once in gelation solution and 4 μL of gelation solution was added to each hole of the gelation chamber. The coverslip was placed on the gelation chamber with the cells facing down and a second 18 mm square coverslip was put on top. Clamps were used to press down the coverslip during gelation to obtain thinner gels (Figure S3D, steps 5-8). The assembled gelation chamber was incubated for 1 hour at 37°C. Gels were removed from the gelation chamber by adding a drop of PBS onto the gels before gently moving them from the Microscopy slide/coverslip into 2 mL Eppendorf tubes using a spatula. Gels were rinsed twice in PBS. After piercing the cap of the Eppendorf tube with a syringe to prevent pressure build-up, non-proteolytic disruption was performed by incubating the gels in disruption buffer (5% SDS (Sigma, 436143-25G), 200 mM NaCl (Sigma, S7279-25G), 50 mM Tris pH 8 (Invitrogen, AM9855G)) O/N at 95°C on a heating block. After disruption, gels were rinsed once in 0.4 M NaCl, followed by two 30 minutes washes in PBS with gentle rocking. Gels were stained with 20 μg/ml DAPI and 20 μg/ml NHS-ATTO 488 or NHS-ATTO 594 in PBS for 1 hour at 37°C. After staining, gels were rinsed once in PBS and transferred to a container with large volumes of Milli-Q water. Gels were expanded for 2x 30 minutes in MilliQ-water with gentle rocking. The water was exchanged again, and gels were left to expand O/N with gentle rocking. Unless mentioned otherwise, all steps were performed at RT. Adapted from (Damstra et al., 2022). The hydrated gels were transferred into a 30 mm imaging dish (Ibidi, 81156) and imaged on a CSU-W1 spinning disk confocal (Evident/Olympus), mounted on a IX83 stand and equipped with diode lasers and Hamamatsu ORCA-Fusion sCMOS cameras. The images were acquired with a 30x UPL S APO 1.05 NA silicon objective (Olympus) and cellSens Dimension software (Evident Scientific, version 4.3.1).

For the experiment shown in Figure S1A: 0.1% Bis was used for gelation. Anchoring was performed overnight in 0.1 mg/ml AcX. Denaturation was performed for 3h at 80°C. These conditions were used to achieve 4x expansion. For any other experiments, the procedure described above was used to allow for 10x expansion.

### Drying of gels onto membrane slides

Fully expanded gels were trimmed with a scalpel and placed on a steel frame slide with PPS-membrane (Leica, 11600294). Excess water was removed with a tissue paper. The slide was incubated at 37°C for 3 hours to dry the gels. Membrane slides were stored at 4°C in a desiccator in the dark until laser dissection.

Widefield images of dried samples (and corresponding non-expanded controls) were captured using the Zeiss Imager.M2 with an EC Plan-NEOFLUAR 20x/0.5 (air) objective and ZEISS ZEN 3.9 Software.

### Laser Microdissection

Laser Microdissection was performed using a LMD7 (Leica Microsystems) equipped with an Ultra LMT350 stage, and a 12-bit K3M monochrome camera. A 349 nm diode-pumped solid-state laser was used for microdissection, and the samples were cut using with a HC PL FLUOTAR 10x/0,30, a HC PL FLUOTAR L 40x/0.60 CORR XT or a HC PL FLUOTAR L 63x/0,70 CORR XT (Leica, 11506505, 11506208 or 11506222 respectively) and samples were collected into a low bind 96-well plate (Eppendorf, EP0030129512). In the Leica LMD software (version 8.5), ROIs were outlined using the drawing tool and collected using the “Middle Pulse” mode; objectives used and number of ROIs collected were adjusted for different sample types (Table S3). Images pre- or post-cuts were done with either a HC PL FLUOTAR 10x/0,30, a HC PL FLUOTAR L 40x/0.60 CORR XT or a HC PL FLUOTAR L 63x/0,70 CORR XT.

### Sample preparation for LC-MS

To avoid sample loss, all buffers were added to the side of the well, followed by a quick spin. Whenever the sample was transferred, low-bind tips were used. After collection 7μL of lysis buffer (100 mM tetra-ethyl ammonium bicarbonate (TEAB) (Supelco, 18597), 0.02% n-dodecyl-ß-maltoside (DDM) (Sigma, D4641), 10 mM tris(2-carboxyethyl)phosphine (TCEP) (Sigma-Aldrich, 626547), 15 mM 2-chloroacetamidecan (CAA) (Sigma, C0267), were added to the sample in the low bind 96-well plate. The sample was heated to 95°C for 1 hour in a thermal cycler. After cooling of the sample to RT, 1 μL of digestion buffer (60% acetonitrile (ACN) (EGT Chemie AG, RC-ACNMS-2.5L), 100 mM TEAB) was added. Next, the sample was again heated to 75°C for 30 minutes in a thermal cycler. After cooling the sample to RT, 2 μL of trypsin solution (10 ng/μL Trypsin (Promega, V5113), 50 mM TEAB, 0.01% DDM) was added and the sample was incubated for 12 hours at 37°C. The reaction was stopped by adding 1 μL of 5% trifluoracetic acid (TFA) (Thermo Fisher, 28904), and the sample was dried in a speedvac (Labconco, centrivap).

Finally, the peptides were cleaned by solid phase extraction using EvoTip Pure stage tips (EVOSEP, EV2020). The dried samples were heated to 37°C, resuspended in 10 μL resuspension buffer (0.1% TFA) using sonication for 5 minutes in a waterbath. The sample were loaded onto EvoTips by centrifugation following the vendor’s protocol. Peptides were eluted with 20 ul elution buffer (40% ACN, 0.1% formic acid (FA)) directly into a low bind 96-well plate, dried under vacuum and stored at -20 °C until further use.

### LC-MS analysis

Peptides were resuspended in 5 μL 0.1% formic acid and the whole sample used for LC–MS/MS analysis using a timsTOF Ultra 2 Mass Spectrometer (Bruker) fitted with a Vanquish Neo HPLC (Thermo Fisher Scientific). The column heater was set to 60°C. Peptides were resolved using a RP-HPLC column (100μm × 30cm) packed in-house with C18 resin (ReproSil Saphir 100 C18, 1.5 μm resin; Dr. Maisch GmbH) at a flow rate of 0.4 μL min^-1^ at 60°C. A linear gradient of buffer B (80% acetonitrile, 0.1% formic acid in water) ranging from 2% to 25% over 25 minutes and from 25% to 35% over 5 minutes was used for peptide separation. Buffer A was 0.1% formic acid in water. The mass spectrometer was operated in positive DIA-PASEF mode. MS spectra were acquired from 100 to 1700 m/z. For MS/MS, we used eight MS/MS ramps and 24 MS/MS windows covering a mass-to-charge ratio range of 400–1000 m/z. This resulted in 25 Th wide MS windows and an ion mobility range from 0.64 to 1.37 V -s/cm². The mass spectrometer was operated in normal sensitivity mode, expected cycle time was 0.95 seconds, accumulation and ramp time was set to 100 ms. The following source settings were applied: 1600V Capillary Voltage and 3 l/min Dry Gas flow at 200°C. The collision energy was linearly ramped between 20 eV at 0.6 V -s/cm² and 59 eV at 1.6 V -s/cm².

## Data analysis

The acquired raw files were searched using SpectroNaut (Biognosys, Schlieren, Switzerland, v19.0, default settings) against a human database (consisting of 20,360 protein sequences downloaded from Uniprot on 2022/02/22) and 393 commonly observed contaminants using the following search criteria: full tryptic specificity was required (cleavage after lysine or arginine residues, unless followed by proline); 3 missed cleavages were allowed; carbamidomethylation (C) was set as fixed modification; oxidation (M), N-acetlyation (N-term) were applied as variable modifications. The raw quantitative data was further statically analyzed using ProteoFlux (v1.8.3, Mollet, D., & Schmidt, A. ProteoFlux (Version 1.8.5) [Computer software]. https://doi.org/10.5281/zenodo.18640999) and Prism (v10.3.1, GraphPad). ProteoFlux is an open-source and reproducible workflow for quantitative proteomics data analysis. Prior to quantification, precursor intensities were filtered based on run-level posterior error probability (PEP) and q-value thresholds (≤ 0.01), and very low-intensity signals (below a threshold of 2) were treated as left-censored measurements. Precursor-level intensities exported from Spectronaut were aggregated to peptide-level quantities and subsequently summarized to protein abundances using a simple summarization-based approach, followed by median-based sample normalization and statistical analysis using linear models with empirical Bayes moderation. Exploratory data analysis included principal component analysis (PCA) of normalized protein intensities after left-censored missing value imputation and visualization of protein identification overlaps between conditions using UpSet plots. For differential proteome analysis, the raw quantitative data was filtered as follows: only proteins quantified with at least two unique peptides and in at least three replicates in one condition per contrast were considered. Subcellular location annotations (GO, cellular compartment) for all human proteins were downloaded from www.uniprot.org (06/02/2026) (Table S6). Functional enrichment analysis was carried out using https://string-db.org/ (Szklarczyk et al., 2025) and the “Proteins with Values/Ranks” search scheme with default settings. Here, all protein accession numbers with their corresponding ratios of a specific contrast were imported. Only enriched terms with a Benjamini-Hochberg corrected FDR rate of 0.01% were considered (Table S7, S11 and S12). The specificity of a specific contrast was determined by dividing the number significant protein hits (ratio >0, adjusted p-value <0.05) with correct subcellular GO annotation to all significant hits. Along these lines, the specificity of published methods was determined in the same way, using the list of all proteins assigned to the corresponding subcellular location as input. Intensity-Based Absolute Quantification (iBAQ) values were obtained directly from SpectroNaut to calculate the total and GO annotated protein mass within the samples.

## Data availability

All data are included in the main text or supplementary. All mass spectrometry proteomics data associated with this manuscript have been deposited to the ProteomicsXchange consortium via MassIVE (https://massive.ucsd.edu) with the accession number MSV000101122/PXD075606.

## Author Contributions

ASpa, CAF, ASch, AF and OB conceived the project. CAF, AF and ASch acquired the data. ASch, CAF, ASpa and AF analyzed and interpreted the data. CAF, ASpa, and ASch wrote the manuscript. All authors commented on the manuscript.

## Acknowledgements

This work was a collaboration between the Spang Lab, the Imaging Core Facility and the Proteomics Core Facility (all Biozentrum, Basel). The authors thank Dariush Mollet for his valuable assistance with data analysis, preparation of figures, and contributions to the methods section. We also thank the Hondele lab (Biozentrum, Basel) for providing antibodies against NCL and SON and the Spiess lab (former Biozentrum, Basel) for sharing the HeLa CCL2 cell line. Finally, we thank Paul Guichard and Virginie Hamel’s lab (Université de Genève) and the FAME facility (FMI, Basel), especially Sabine Reither for productive discussions on expansion protocol optimization. This work was supported by the University of Basel and the Swiss National Science Foundation (310030_197779, 310030_219513, 320030-231859) to A. Spang.

---

**Figure S1**

**(A)** Number of identified proteins for different input cell numbers. Data are mean ± SD: 50 cells: 2964; 10 cells: 2374; 5 cells: 2417; 1 cell: 982.5 ± 220.6 (n = 1 for 50, 10, and 5 cells; n = 6 for 1 cell; N = 1 experiment). **(B)** Hydrogels before and after drying for 3 hours at 37°C onto PPS-membrane slides. **(C)** Number of peptides identified either terminating with lysine (K) or arginine (R) across all samples shown in Figure 2F. **(D)** Volcano plot of pairwise proteomic comparison between nuclei and whole cells. Red dots represent proteins annotated as nuclear according to GO. **(E)** Volcano plot of pairwise proteomic comparison between nuclei and CellsMinusNuclei (same data as Figure 3B). Red dots represent proteins annotated as mitochondrial and not nuclear according to GO. **(F)** Schematic illustrating the SPEx “standard” approach and the SPEx “subtraction” approach. In the standard approach, the proteome of the organelle of interest (e.g., nuclei) is compared to the proteome of whole control cells from which the organelle was removed. In the subtraction approach, the proteome of a whole cell is compared to the proteome of a cell lacking the organelle of interest. In both approaches, proteins localized to the organelle of interest are enriched, and absent proteins are depleted. Created in BioRender.com. **(G)** Venn diagram depicting the overlap of identified nuclear proteins between the standard SPEx (Figure 3B) and subtraction SPEx approach (Figure 3G). **(H)** Bar chart illustrating the number of proteins in the overlap shown in (G) with (Yes) and without (No) nuclear GO annotation. **(I)** Bar plot comparing the number of proteins assigned to the nucleus (blue) in our SPEx dataset (this study, Figure 3B) with other published spatial proteomics datasets (Geladaki et al., 2019; Hein et al., 2025; Orre et al., 2019). Proteins overlapping with nuclear GO annotation are shown in red. **(J)** Specificity of nuclear GO annotation in the different proteomics datasets, determined from (I).

**Figure S2**

**(A)** Volcano plot of pairwise proteomic comparison between nucleoli and NucleiMinusNucleoli. The dotted square indicates proteins significantly enriched in nucleoli (adjusted *P* < 0.05, log₂FC > 0). Red dots represent proteins annotated to localize to the nucleolus according to GO. **(B)** Number of proteins significantly enriched in nucleoli in (A, dotted square) that are annotated as nucleolar (Yes) or not (No) according to GO. **(C)** Protein mass composition of nucleolus samples. Red bar indicates the percentage of protein mass contributed by proteins annotated as nucleolar according to Gene Ontology (GO) to the total protein mass based on iBAQ values. **(D)** A-431 cells immunostained for RBM5 and tubulin. Image adapted from Protein Atlas (www.proteinatlas.org). Scale bar, 10 µm. **(E)** A-431 cells immunostained for SNW1 and tubulin. Image adapted from Protein Atlas (www.proteinatlas.org). Scale bar, 10 µm. **(F)** Bar plot comparing the number of proteins assigned to the nucleolus (blue) in our SPEx dataset (this study, Figure 4A) with the published proteomics dataset (Orre et al., 2019). Proteins overlapping with nucleolar GO annotation are shown in red. **(G)** Specificity of nucleolar GO annotation in the different proteomics datasets, determined from (F).

**Figure S3**

**(A)** Protein mass composition of Golgi samples. Red bar indicates the percentage of protein mass contributed by proteins annotated as Golgi and/or extracellular according to Gene Ontology (GO) to the total protein mass based on iBAQ values. **(B)** Volcano plot of pairwise proteomic comparison between WholeCells and CellsMinusGolgi. The dotted square indicates proteins significantly enriched (adjusted *P* < 0.05, log₂FC > 0) in the WholeCells. Red dots represent proteins annotated to localize to the Golgi and/or to be extracellular according to GO. **(C)** Number of proteins significantly enriched in the Golgi in (B) that are annotated as Golgi and/or extracellular (Yes) or not (No) according to GO. **(D)** Gelation chamber assembly and sample mounting workflow. (1) Two holes of 3-4 mm diameter (spaced apart by ∼3mm) were punched into a microscopy-slide-sized piece of parafilm using a tissue puncher. (2) The parafilm was placed on a glass microscopy slide (paper side up) and adhered by repeatedly streaking over it with a second microscopy slide. (3) The paper on the parafilm was removed using tweezers. (4) A non-adhesive plastic foil was put over the parafilm and the streaking with a second microscopy slide was repeated until the parafilm was perfectly adhered to the microscopy slide. (5) 4 μL of gelation solution was added to each hole of the gelation chamber (6) The 13 mm round coverslip containing the cells was rinsed in gelation solution and placed on the gelation chamber with the cells facing down. (7) An empty 18 mm square coverslip was put on top of the round coverslip containing the sample (8) Clamps were used to press down the coverslips during gelation to obtain thinner gels.

**Supplementary video 1: Subcellular laser microdissection of different samples**

Example movies of laser microdissection for all samples shown in Figure 2E. The movies are shown at 3x speed.

