## Supplementary figures and images for "SPEx: Compartment-Resolved Proteomics via Expansion Microscopy–Guided Microdissection"

### suppl_Figs.pdf

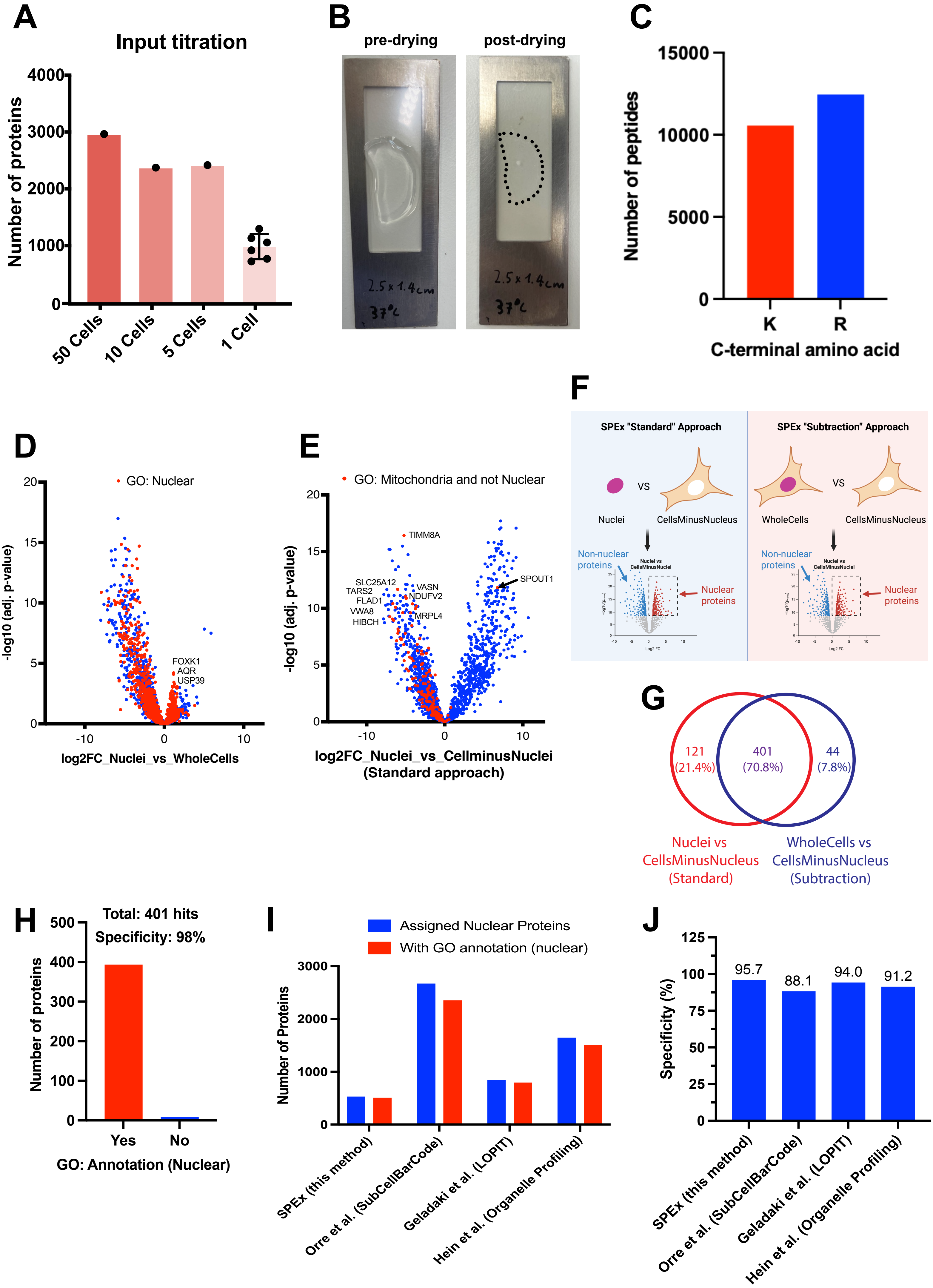

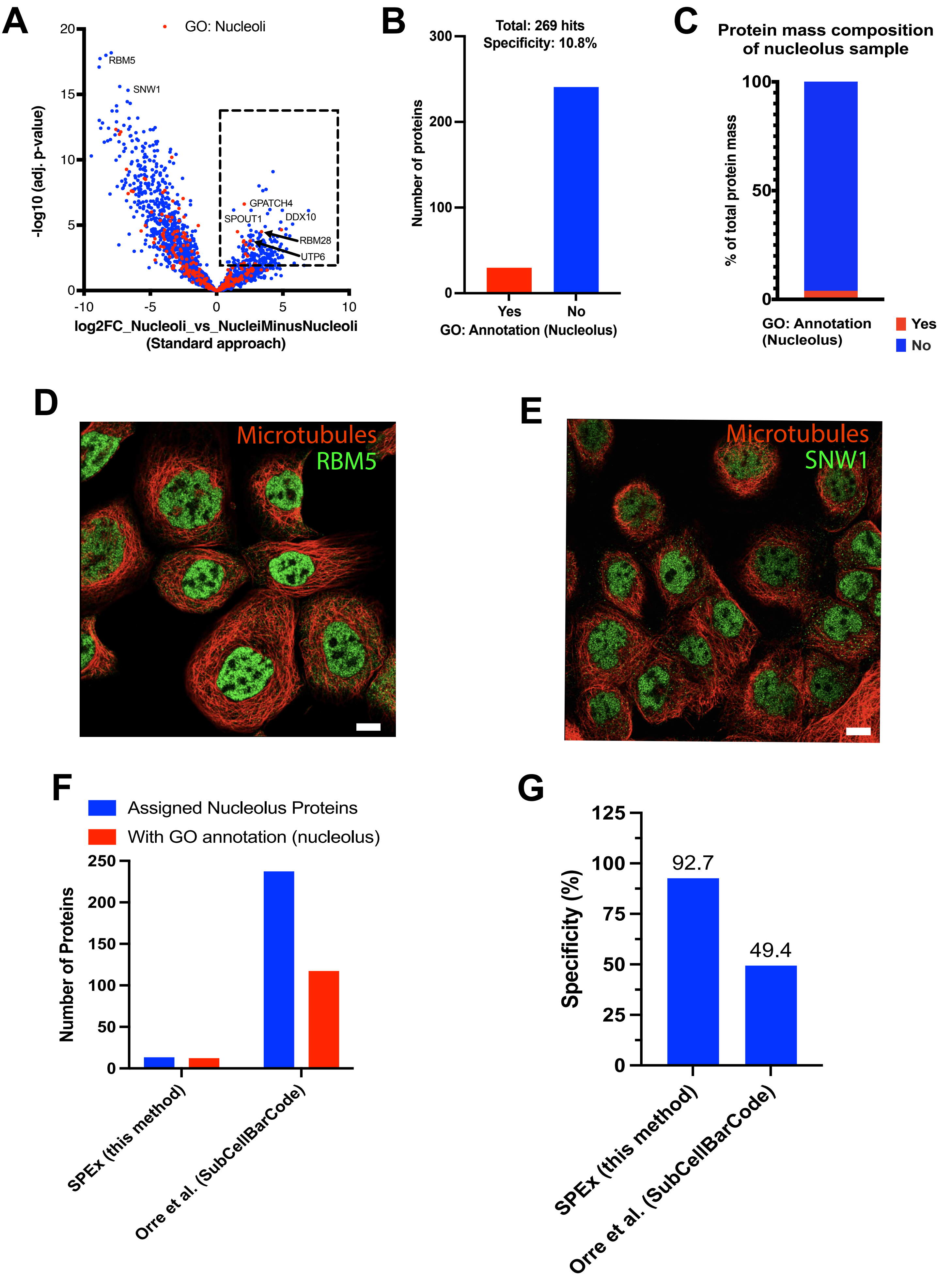

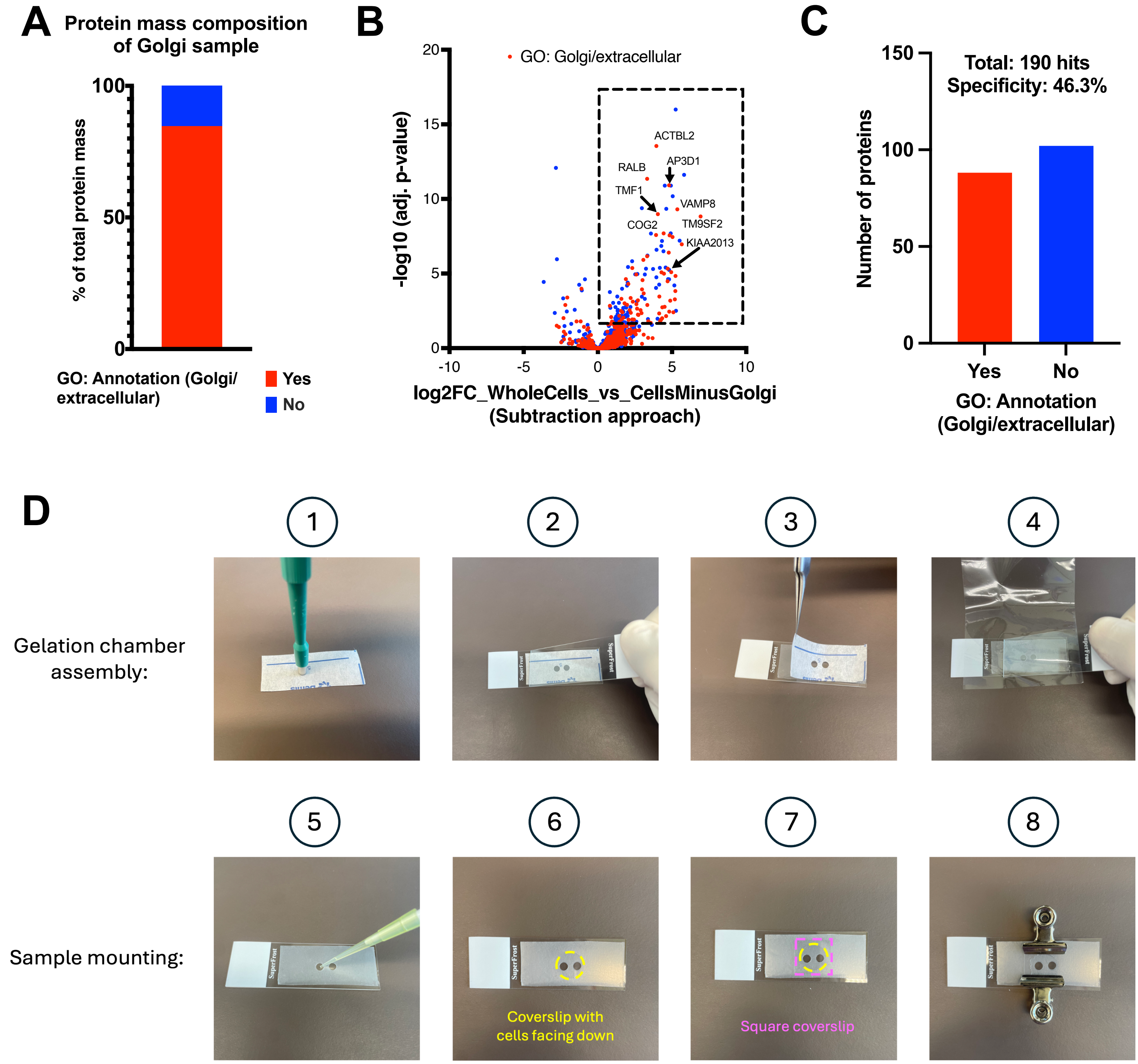
